# Multiplex Enrichment and Detection of Rare KRAS Mutations in Liquid Biopsy Samples using Digital Droplet Pre-Amplification

**DOI:** 10.1101/298299

**Authors:** Erica D. Pratt, Robert W. Cowan, Sara L. Manning, Edmund Qiao, Heather Cameron, Kara Schradle, Diane Simeone, David B. Zhen

## Abstract

Oncology research is increasingly incorporating molecular detection of circulating tumor DNA (ctDNA) as a tool for cancer surveillance and early detection. However, non-invasive monitoring of conditions with low tumor burden remains challenging, as the diagnostic sensitivity of most ctDNA assays is inversely correlated with total DNA concentration and ctDNA abundance. Here we present the Multiplex Enrichment using Droplet Pre-Amplification (MED-Amp) method, which com-bines single-molecule emulsification and short-round PCR preamplification with digital droplet PCR (ddPCR) detection of mutant DNA template. The MED-Amp assay increased mutant signal by over 50-fold with minimal distortion in allelic frequency. We demonstrate detection of as few as 3 mutant copies in wild-type DNA concentrations ranging from 5 to 50ng. The MED-Amp assay successfully detected KRAS mutant ctDNA in 86% plasma samples obtained from patients with metastatic pancreatic ductal adenocarcinoma. This assay for high-sensitivity rare variant detection is appropriate for liquid biopsy samples, or other limited clinical biospecimens

## Introduction

Prior research has shown that digital droplet PCR (ddPCR) has superior accuracy to traditional quantitative PCR (qPCR) for detection of rare genetic mutations. ddPCR also outperforms qPCR in biological fluids containing natural PCR inhibitors ^1^, more accurately distinguishes between rare events and false-positives ^2^, and minimizes measurement variation by partitioning DNA template into thousands to millions of discrete reaction volumes. Each droplet becomes an individual PCR reaction event, increasing target signal-to-noise ratio, and theoretically enabling detection of mutations with allelic frequencies lower than 0.1%. Due to its sensitivity and cost-effectiveness, ddPCR is an attractive alternative to next-generation sequencing for targeted detection of rare mutations. Thus, ddPCR is increasingly used in cancer research for high-sensitivity molecular detection of circulating tumor DNA (ctDNA) for non-invasive detection and monitoring of disease. Circulating tumor DNA is released into the circulation from tumor cells and is a non-invasive prognostic indicator of survival ^3^. However, the majority of cell-free DNA (cfDNA) in the circulation is wild-type and may contain as little as one mutant DNA fragment per milliliter of plasma ^3, 4, 5^. Prior work has demonstrated ddPCR is a powerful tool for ctDNA detection and serial monitoring for patients with advanced disease and high tumor burden ^6, 7^. However, diagnostic sensitivity is inversely correlated with total cfDNA concentration, limiting detection in samples where either wild-type or mutant DNA concentrations are low ^5, 8, 9^, such as serial monitoring of minimal residual disease. An especially salient example is pancreatic cancer, where over 90% of primary tumors contain a *KRAS* mutation ^10^. Yet in recent studies of patients with confirmed *KRAS* mutant positive tumors, only 35-43% of plasma samples also tested positive via ddPCR ^5, 11^.

To further increase assay sensitivity for detection of low DNA input and low target abundance samples, several groups have performed short cycles of conventional PCR with a high-fidelity polymerase to increase the starting concentration of nucleic acids prior to downstream detection. A recent study showed that preamplification of template increased true-positive ddPCR signal from 15-to 27-fold with only nine cycles of PCR with only a 1.5-to 9-fold increase in false positives. This preamplification method enabled detection of 0.05% mutant *KRAS* ctDNA in 50 ng of wild-type DNA ^12^. Furthermore, multiplexed detection of cancer-specific mutations was possible with total cfDNA inputs as low as 9 ng. Similarly, fifteen cycles of preamplification prior to next-generation sequencing resulted in detection of mutant transcripts at 0.63% allelic frequency with DNA inputs as low as 2 ng ^13^. However, conventional PCR is sensitive to amplification bias based on DNA fragment size, resulting in distortion of the original allelic fraction. Circulating tumor DNA is highly fragmented, exists at very low concentrations, and is systematically shorter than wild-type cfDNA ^14^, making it particularly susceptible to these biases. Additionally, prior research has shown a 10-20% reduction in PCR amplification efficiency for mutation-containing DNA fragments in common tumor-specific genes, such as *TP53* and *KRAS* ^*15*^.

Here we describe a new assay, <u>M</u>ultiplex <u>E</u>nrichment using <u>D</u>roplet Pre-<u>Amp</u>lification (MED-Amp), which addresses the limitations described above. This method combines single-molecule DNA emulsification in picoliter volume droplets with nine rounds of preamplification using a high-fidelity polymerase, followed by ddPCR detection of template. The MED-Amp assay has the capacity to identify single mutant DNA fragments from wild-type using cfDNA inputs as low as 5 ng. We show that the MED-Amp method generates linear DNA template amplification, enabling back-calculation of original ctDNA concentrations. Finally, we piloted our assay with plasma samples from patients with metastatic pancreatic ductal adenocarcinoma (PDA) for multiplexed detection of the four most common *KRAS* codon 12 mutations in pancreatic cancer (p.G12C, p.G12D, p.G12R, p.G12V) ^10^.

## Results

### Multiplex KRAS mutation targeting

Our multiplex mutation detection method consisted of DNA template emulsification and PCR amplification in droplets prior to de-emulsification and repartitioning for analysis using standard TaqMan® chemistry (Figure 1). In our assay, DNA template and high-fidelity polymerase master mix were loaded on the RainDance Source digital PCR system for single-molecule partitioning in 5 picoliter volume droplets. After nine rounds of PCR amplification, the resulting PCR product was de-emulsified, purified, and repartitioned with TaqMan® Genotyping Master Mix combined with TaqMan® probes for *KRAS* detection.

**Figure 1.**
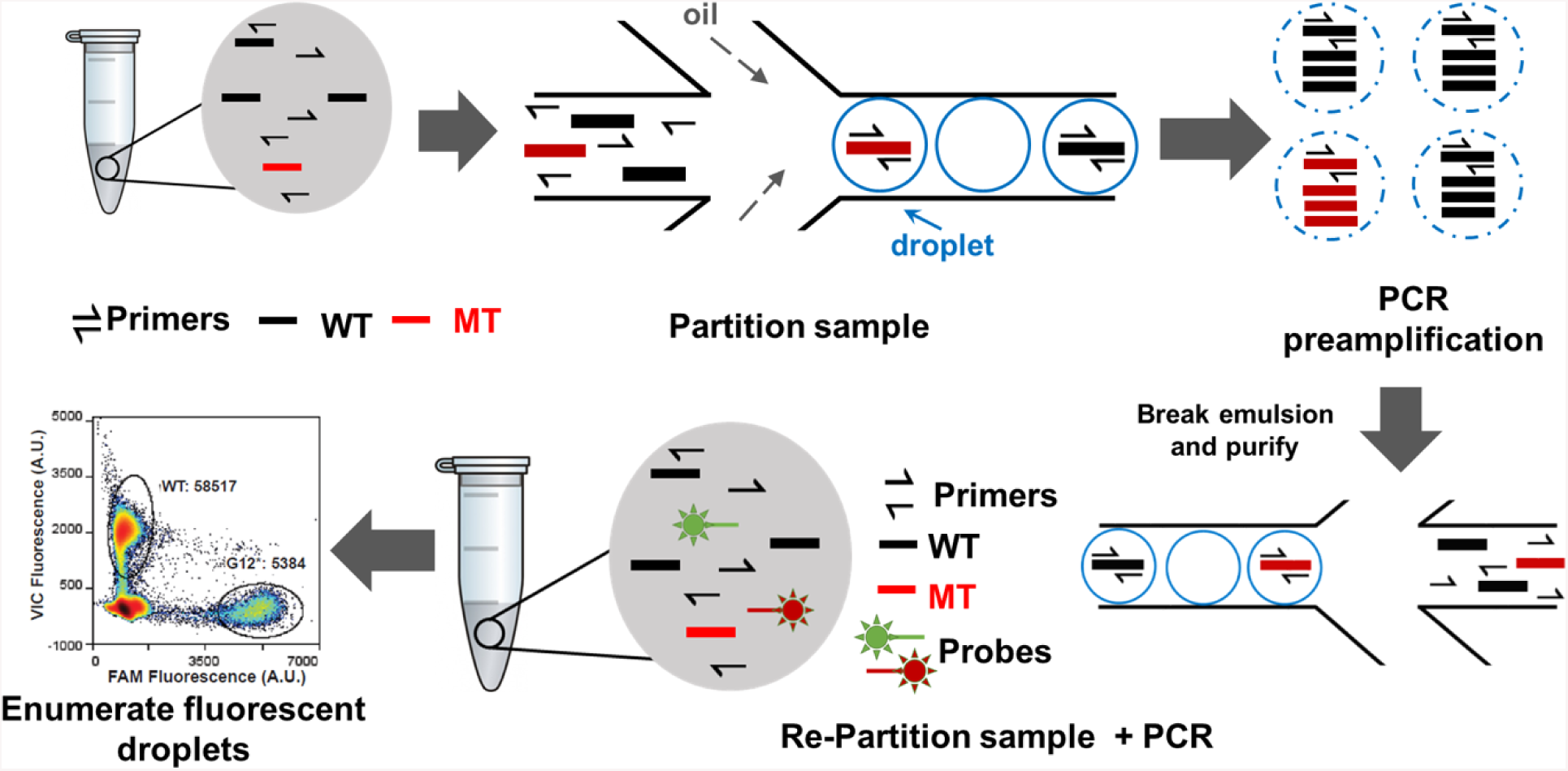
Experimental workflow for MED-Amp. Master mix containing DNA template, primers, and PCR reagents are partitioned into 5 pL droplets and undergo 9 rounds of preamplification. The emulsion is broken and the PCR product is purified using a PCR cleanup kit, then re-partitioned with primers and TaqMan® probes for digital PCR detection.

We designed a multiplexed KRAS panel, combining TaqMan® probes targeting wild-type sequence, and the most common KRAS codon 12 mutations in PDA (p.G12C, p.G12D, p.G12R, p.G12V) ^10^. Based on biorepository data (ICGC, QCMG, and TCGA) encompassing over 700 patients, these four *KRAS* mutations are present in 84% of PDA samples. Previous studies have shown ctDNA is heavily fragmented, averaging 160 bp in length ^14, x16^. Therefore, we designed a *KRAS* primer set producing a 95 bp amplicon flanking the codon 12 mutation sites of interest (SI Table 1). To maximize assay sensitivity, a FAM reporter was used for all four *KRAS* mutant probes and concentrations were optimized such that all FAM-positive events clustered into a single gate. FAM-positive events would then achieve maximum separation from the much stronger wild-type VIC signal. We confirmed successful probe multiplexing into a single G12 mutation gate (labeled G12*) using a mixture of DNA from various pancreatic cell lines encompassing all four *KRAS* mutations (data not shown).

**Table 1.** Patient Characteristics

| Sample ID | Age | Sex | Pathology | Tumor Location | Tumor Size (cm) | Prior Therapy | DNA Input (ng) | AF (%) | WT Droplets | MT Droplets |
| --- | --- | --- | --- | --- | --- | --- | --- | --- | --- | --- |
| S03 | 46 | M | Control | -- | -- | -- | 4.0 | 13.5 | 25612 | 3989 |
| S04 | 55 | M | Control | -- | -- | -- | 2.4 | 0.03 | 277324 | 83 |
| S05 | 47 | M | Control | -- | -- | -- | 18.2 | 0.20 | 12731 | 23 |
| S06 | 58 | M | Control | -- | -- | -- | 2.5 | 0.01 | 33993 | 5 |
| S07 | 53 | F | Control | -- | -- | -- | 3.4 | 0.01 | 32744 | 4 |
| S08 | 49 | F | Control | -- | -- | -- | 1.5 | 0.15 | 4661 | 7 |
| S09 | 51 | M | Control | -- | -- | -- | 1.2 | 0.06 | 9465 | 6 |
| S10 | 83 | F | Control | -- | -- | -- | 1.4 | 0.05 | 13376 | 7 |
| S11 | 74 | M | PDA | Head | 2.6 | None | 10.2 | 5.4 | 87900 | 5016 |
| S12 | 58 | F | PDA | Body | 8.8 | None | 2.8 | 8 | 57904 | 5023 |
| S13 | 55 | F | PDA | Body/Tail | 5.5 | None | 4.1 | 13.4 | 39260 | 6097 |
| S14 | 75 | F | PDA | Head | 9.1 | None | 17.0 | 0 | 219011 | 5 |
| S15 | 72 | F | PDA | Tail | 5.4 | None | 12.0 | 0.04 | 76245 | 34 |
| S16 | 62 | F | PDA | Head | 3 | Chemotherapy,<br>Radiation | 3.6 | 0.24 | 13712 | 33 |
| S17 | 66 | M | PDA | Head | 8 | Chemotherapy | 13.0 | 0.14 | 124335 | 173 |

\*Shaded rows indicate positive KRAS detection by ddPCR

### Detection of mutant alleles using droplet-based preamplification

To recapitulate circulating tumor DNA concentrations seen in plasma samples, we created twelve artificial dilutions of a *KRAS* gene-specific multiplex reference standard containing a characterized *KRAS* p.G12D mutation at 16.7% allelic frequency. Dilutions were created by mixing the reference standard with DNA from the BxPC3 human PDA cell line, which is homozygous for wild-type *KRAS* at codon 12. *KRAS* mutant copy number ranged from 3 to 272 in a total wild-type background of 1,426 to 14,257 (5-50ng) genomic equivalents, resulting in allelic frequencies spanning 0.02% to 8.71%. Each allelic frequency was replicated at least fourteen times.

We successfully identified *KRAS* p.G12D mutations in 87% of samples tested (Figure 2a). There was high concordance between measured *KRAS* mutant fraction post-preamplification and nominal expected fraction across the allelic frequencies tested (Figure 2b). Post-preamplification FAM signal was linearly correlated with input mutant DNA copy number as described by a generalized least squares regression model with an exponential variance function to correct for heteroscedasticity (p<0.001). The linearity of amplification across a two order-of-magnitude change in mutant template copy number allows for straightforward back-calculation of original sample VAF.

**Figure 2.**
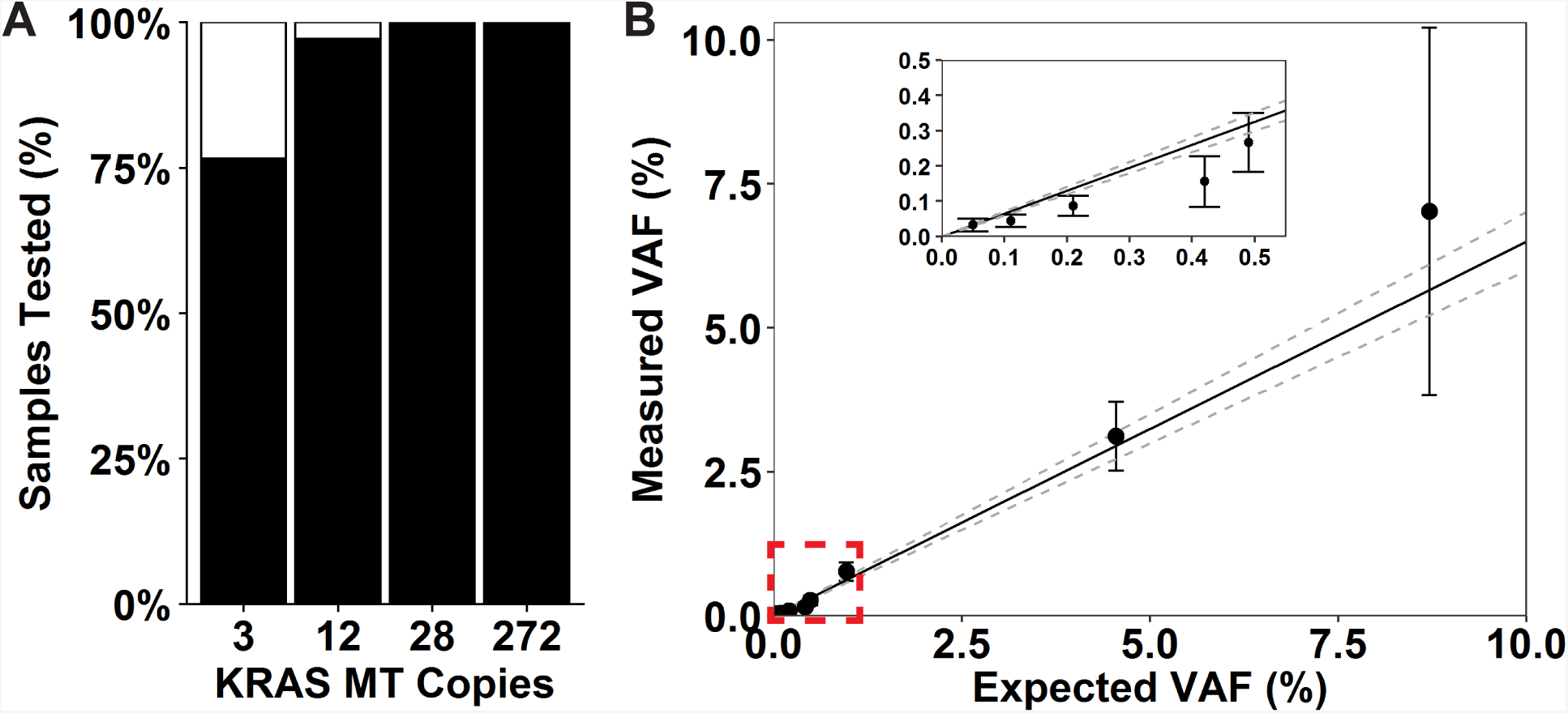
A) MED-Amp detection sensitivity as a function of input mutant *KRAS* copy number. B) Nominal expected frequency versus measured allelic frequency (VAF). Best fit generated using generalized least squares regression model with heteroscedasticity correction (y=0+0.650*x). Inset shows < 0.5% allelic frequency data for clarity.

To assess overall assay performance, we then compared the number of mutant *KRAS* droplets detected via ddPCR with and without preamplification. Amplification ratio, defined as number of *KRAS* mutant droplets measured by ddPCR divided by number of spiked *KRAS* mutant copies, was used to characterize preamplification efficiency. Each allelic frequency tested was assayed in triplicate. Emulsified preamplification increased mutant *KRAS* signal an average of 50-fold with only an 8-fold increase in false positives from non-template or wild-type *KRAS* controls (Figure 3a & 3b), consistent with other preamplification-based techniques ^12^. While preamplification resulted in a mean 50-fold increase in signal, this falls well short of the over 500-fold increase expected from nine cycles of preamplification. These data are consistent with prior reports of suppressed amplification using a variety of high-fidelity polymerases for preamplification ^12, 13^. Amplification was uniform across a two order of magnitude change in mutant *KRAS* copy number and allelic fraction (Figure 3c). However, unlike other preamplification protocols ^8, 9^, MED-Amp had no dependence on total DNA input (SI Table 2). These data show emulsified preamplification may improve the sensitivity of detection of variant alleles at extremely low concentrations.

**Figure 3.**
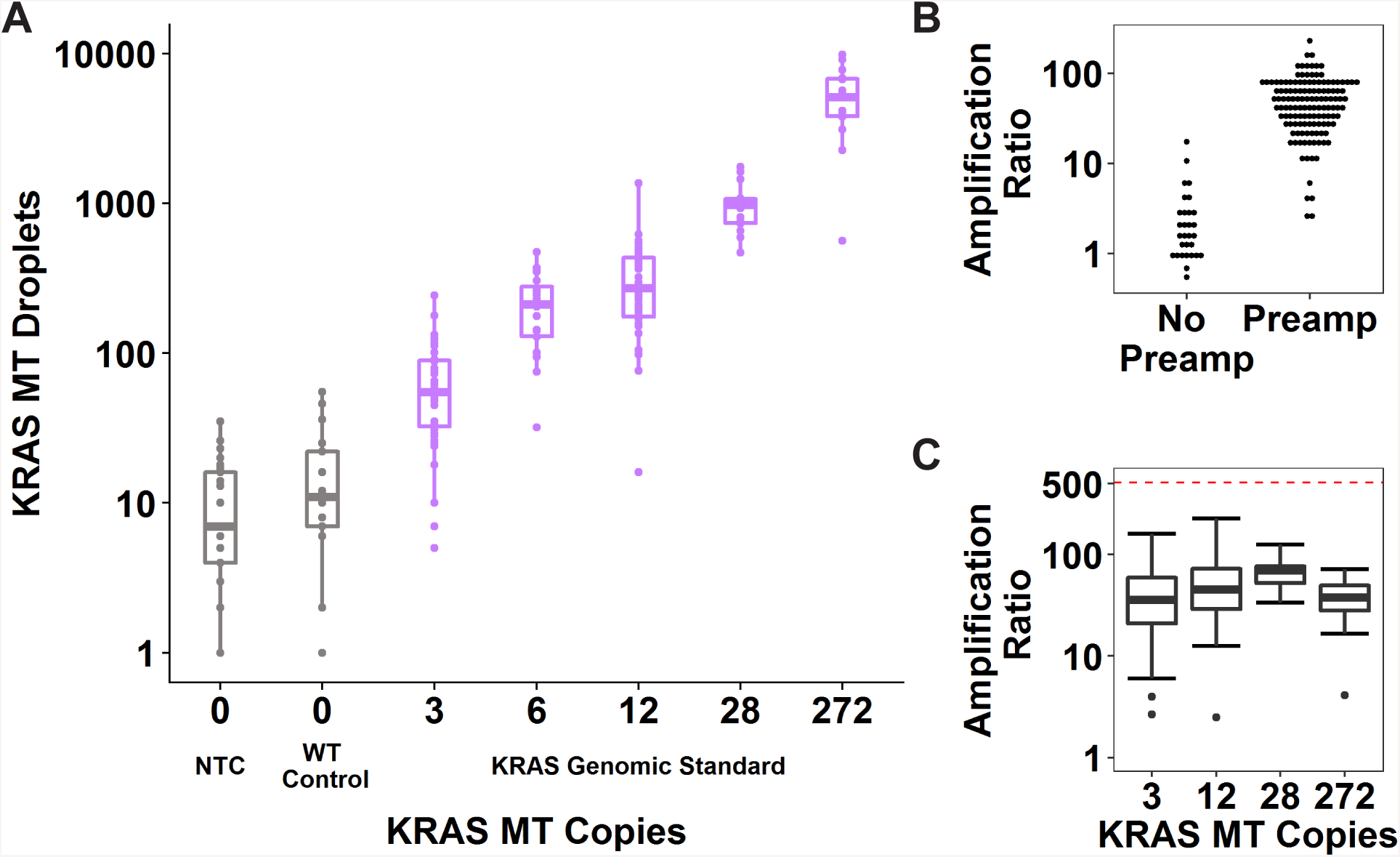
Evaluation of pre-amplification in droplets for detection of KRAS codon 12 mutations. A) Signal amplification as a function of *KRAS* mutant (MT) copy number. B) Average amplification ratio (defined as *KRAS* mutant signal divided by number of spiked *KRAS* mutant copies) of DNA standards containing the *KRAS* p.G12D mutation with and without preamplification. C) Variation in amplification ratio as a function of number of *KRAS* mutant copies present in the DNA sample. Dashed line is the expected amplification efficiency for a perfectly efficient PCR reaction.

### Droplet preamplification improves variant allele detection compared to conventional PCR preamplification

We also compared droplet-versus conventional-PCR preamplification (Figure 4). Overall amplification ratio was not correlated with template emulsification (p = 0.38). However, droplet amplification was two-fold more efficient than conventional amplification as the allelic frequency dropped below 0.2%, near the limit of detection of most ddPCR assays (p = 0.03) (Figure 4a). Importantly, conventional preamplification consistently underestimated mutant *KRAS* template allelic frequency, and was on average 59% lower than droplet preamplification (Figure 4b & 4c). Therefore, droplet preamplification was further developed for highly sensitive and specific detection of rare variants.

**Figure 4.**
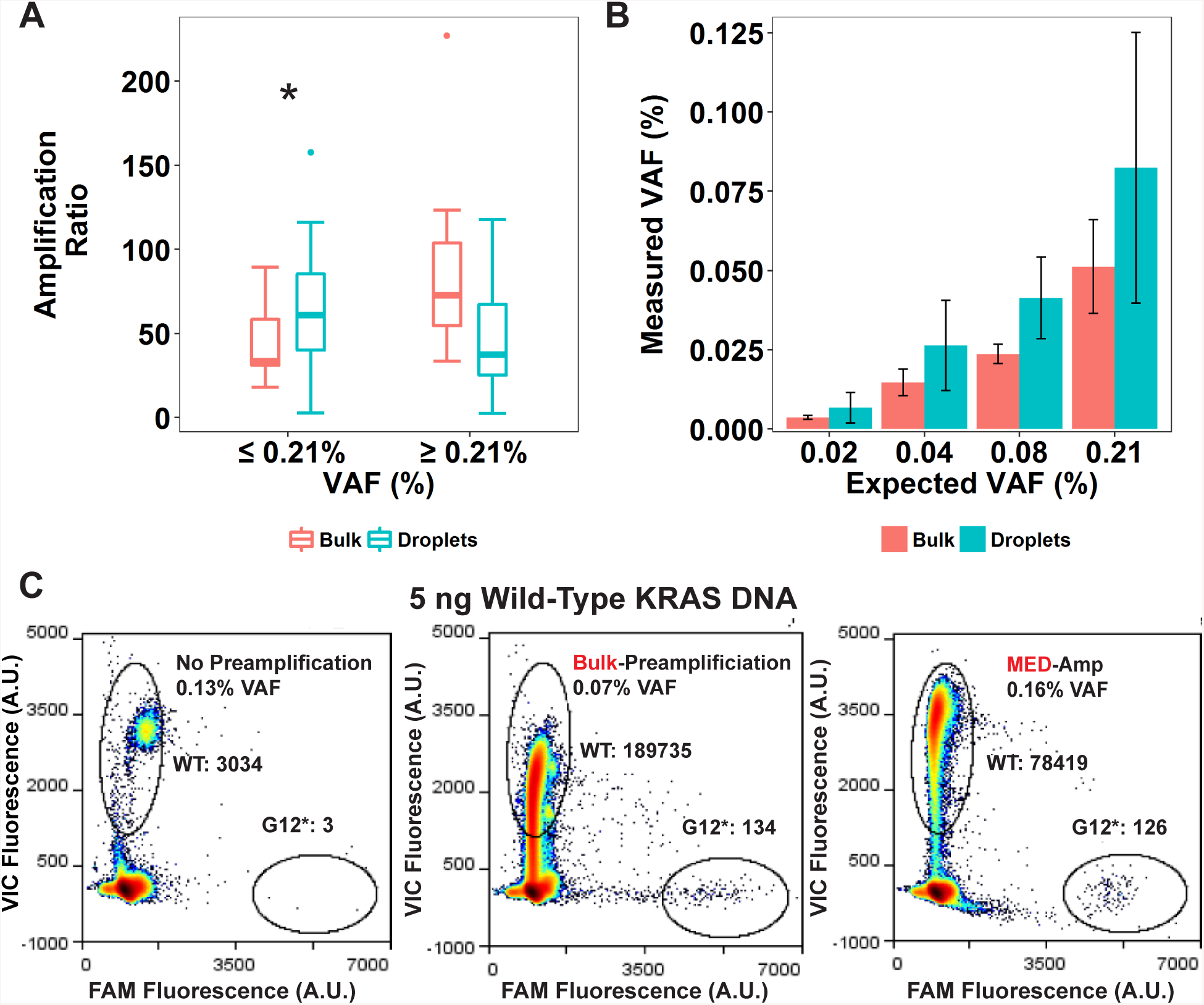
Comparison of ddPCR amplification efficiency and measurement fidelity in bulk preamplification versus MED-Amp. A) Amplification ratio dependence on relative abundance of *KRAS* mutant DNA. * p = 0.03 B) allelic frequency fidelity for bulk preamplification versus preamplification in droplets. C) Representative ddPCR results for each amplification condition versus a non-preamplification control. Fluorescence intensity is measured in arbitrary units (A.U.)

### Evaluation of high-fidelity polymerase performance

We next compared the performance of three commercially available high-fidelity polymerases, Q5® Hot Start High-Fidelity, PfuUltra II Fusion HS, and Platinum™ SuperFi™, in our assay. Each polymerase was tested at minimum four types at each mutant DNA copy number assayed. While all three polymerases were effective, Q5® led to a 59-fold increase in detected *KRAS* signal compared to PfuUltra II Fusion HS (50-fold) and Platinum™ SuperFi™ (30-fold) (Figure 5a). This trend remained after normalization for amplification variability by input mutant *KRAS* copy number. Q5® and PfuUltra II exhibited higher, more variable, signal amplification, while Platinum™ SuperFi™ signal amplification was lower and more consistent (Figure 5b). We also compared the limit of detection (LOD) for the three polymerases using our assay. All three polymerases produced minimal false-positives in non-template controls: on average seven droplets for Platinum™ SuperFi™, ten for Q5®, and seventeen for PfuUltra II. These data informed limit of detection calculations for each polymerase. We performed a chi-squared goodness-of-fit test and found droplet counts did not fit a Poisson distribution as is commonly assumed in analytical sensitivity measurements ^17, 18^. Therefore, LOD was defined as three standard deviations from the mean false-positive count of each polymerase non-template control ^18^. Based on these LOD calculations, Q5®, Platinum™ SuperFi™ and PfuUltra II detected three mutant *KRAS* molecules with 75%, 70%, and 60% sensitivity and 100% specificity, respectively (SI Table 2). Q5® High-Fidelity polymerase was selected for further assay optimization based on its performance in detecting extremely low amounts of *KRAS* mutations, efficiency in amplification, and specificity.

**Figure 5.**
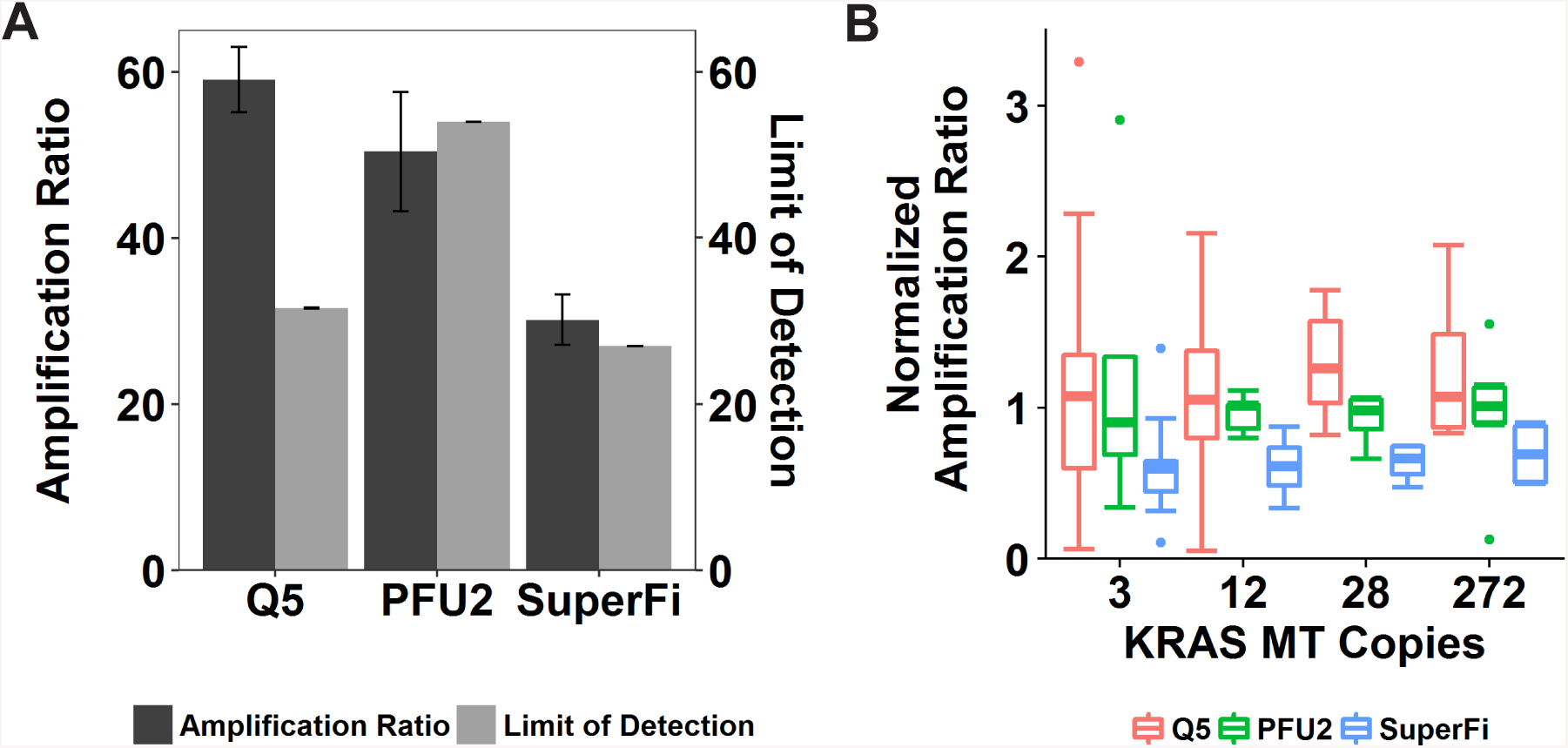
Comparison of High-fidelity polymerases for DNA preamplification. A) Amplification ratio for each high-fidelity polymerase versus its corresponding limit of detection. B) Amplification ratio for each polymerase normalized by mean amplification efficiency per input mutant *KRAS* copy number. C)

### MED-Amp detection of KRAS mutations in plasma

The genomic standards described previously were spiked into 1 mL of plasma from donors without cancer, and the DNA was isolated and assayed as described above. Each copy number input was assayed at minimum in triplicate. Overall assay sensitivity in spiked plasma samples was 67.7% for fewer than 20 mutant DNA copies and 91% for 20 copies and above. Despite the presence of natural PCR inhibitors in plasma, and losses associated with the DNA isolation process, assay sensitivity remained high and compared favorably to detection rates using DNA reference standards alone (Figure 6a).

**Figure 6.**
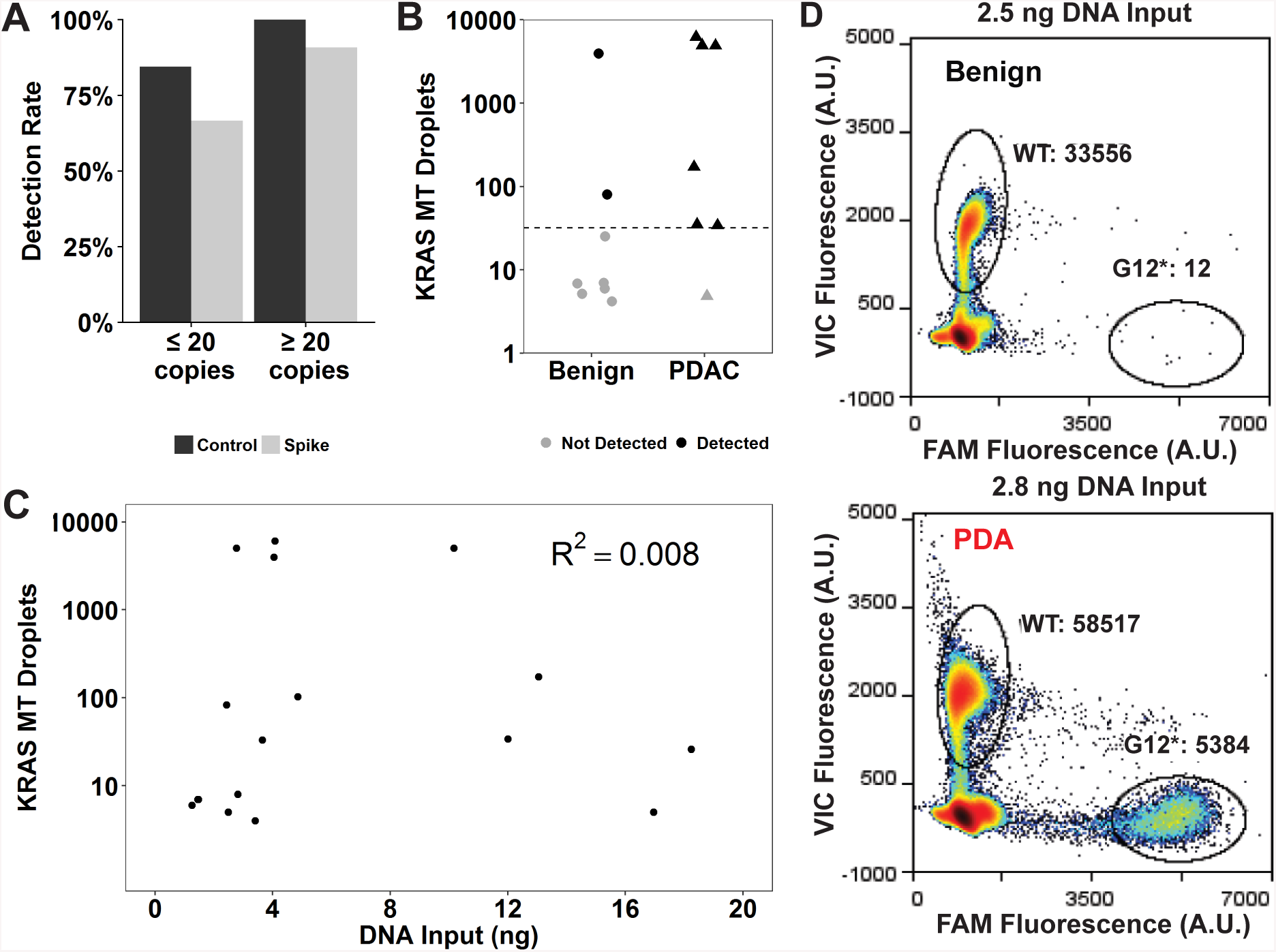
A) MED-Amp detection rate in spiked plasma samples compared to DNA reference standards alone based on input mutant DNA copy number. B) Correlation between ddPCR signal for mutant *KRAS* and total DNA loaded for the PI-Amp assay. C) Mutant *KRAS* signal in plasma from metastatic pancreatic cancer patients versus healthy controls. Dashed line indicates the limit of detection (LOD), defined as three standard deviations from the mean false-positive count. D) Representative ddPCR results for low-input DNA samples (< 5 ng) for a benign control versus a PDA patient.

### MED-Amp detection of KRAS mutations in patient samples

We next applied our assay to PDA patient plasma samples (Figure 6b, Table 1, SI Table 4). We measured *KRAS* mutations in blinded plasma samples from metastatic PDA patients (n = 7), and age-matched non-PDA patient controls (n = 10). Cell-free DNA was isolated from 1-2 mL of plasma from each patient. Final DNA concentrations ranged from 3 ng/mL to 49 ng/mL and were significantly higher in PDA samples compared to controls (p < 0.001, SI Figure 2). The maximum template volume of 8 μL per 25 μL reaction volume was used for each preamplification reaction (1.2 ng to 18 ng total). There was no correlation found between cfDNA concentration and age or sex across groups. Additionally, mutant *KRAS* detection did not correlate with total DNA input (Figure 6c). This is in contrast to prior reports where *KRAS* variant detection was correlated with higher DNA input, which does not itself necessarily correlate with disease activity ^5, 9^.

*KRAS* codon 12 mutations were identified in 6 out of 7 PDA patient samples tested (Figure 6b & 6d), resulting in a diagnostic sensitivity of 86%. Three out of ten control samples also tested positive for mutant *KRAS*, yielding a specificity of 70%. Samples positive for the mutation had no correlation with age for either cohort. Measured allelic frequencies were highly heterogeneous in patient samples, ranging from 0.04% to 13.4%. The average mutant allelic frequency was much higher in PDA samples (3.89%) compared to controls (1.43%, p = 0.33) (SI Figure 2). Based on input DNA concentrations, the total number of mutant DNA fragments present in the tested plasma samples ranged from 1.56 to 195 copies.

We then examined the controls samples which tested *KRAS*-positive. Controls with sufficient remaining plasma, or isolated cfDNA, were re-analyzed using standard ddPCR, and the presence of codon 12 *KRAS* mutations was confirmed (SI Figure 3). Two controls were censored after review of patient records revealed a prior history of cancer, one of which tested mutant *KRAS* positive by MED-Amp. Of the remaining eight patients, six had a history of colorectal polyps. Prior research has identified *KRAS* codon 12 mutations in 15-38% of patients with polyps ^19^. Recent studies have also reported similar false-positive rates in patients with no history of cancer ^20, 21, 22, 23, 24, 25^. These findings suggest somatic *KRAS* mutations may also be a marker of neoplasia or other premalignant conditions.

## Conclusion

Current ddPCR-based genotyping studies have three major limitations: 1) a strong interdependence between total DNA input and assay sensitivity ^5, 8, 9^, 2) low rates of ctDNA detection as compared to matched tumor tissue specimens ^3, 4,5^, and 3) misestimation of ctDNA allelic frequency. cfDNA input in the sub-5 ng range is a disqualifying factor for commercial digital sequencing assays ^26^. In contrast, by using a droplet preamplification step with a high-fidelity polymerase, the MED-Amp method accurately detects low abundance mutations in sub-5 ng DNA input samples. Preamplification in droplets may be a critical step, as Barnard and colleagues reported a strong PCR bias against point mutations within CXGG motifs in *KRAS* codon 12, as well as *TP53* codons 248 and 282, when co-amplified with wild-type sequence ^15^. These data corroborate our observation that conventional PCR preamplification underestimates mutant *KRAS* allelic frequency. Barnard et al. found the observed PCR bias was abated by amplifying wild-type and mutant template in separate reactions, which is analogous to partitioning DNA molecules into individual droplets to serve as compartmentalized PCR reaction vessels. Hence, the emulsification of template prior to PCR leads to more efficient template amplification than conventional PCR preamplification, while also preserving the allelic distribution of the original DNA sample.

The high efficiency of mutant template amplification could explain our reported 86% *KRAS* ctDNA detection rate, which is comparable to methods employing more expensive next-generation sequencing. In contrast, only 35% to 64.7% of late-stage metastatic patients tested positive in other ddPCR-based assays ^4, 27, 28^. Further validation in a larger patient cohort is needed, but our pilot data suggest droplet preamplification could achieve results similar to NGS-based methods and performed at less cost (approximately $200 per sample).

Interestingly, we found that 30% of controls tested positive for *KRAS* mutations. There is a growing body of showing *KRAS* and other tumor biomarkers are present at low levels in individuals with no evidence of malignant disease. These observations have been seen in cfDNA and exosomal DNA, at rates ranging from 5% - 20%, consistent with the results reported here ^19, 21, 23, 25, 22, 24^. The preponderance of studies reporting similar rates of oncogenic mutations using various detection methods, genetic targets, and ranges of DNA inputs suggests further investigation into the prevalence of these mutations in aging populations is warranted. Furthermore, the majority of control samples are obtained at single-time points, and the lack of serial blood draws for analysis limits the ability to determine if the above results represent actual false-positives or early detection of malignant conditions.

While this study focused exclusively on PDA, the method described here could be applied to other tumor types and biospecimens including fine needle aspirates ^29^, as well as archived FFPE or frozen tissue samples. Assay sensitivity remained high, even when applied to frozen samples under extended storage conditions (> 2 years), enabling retrospective analysis of stored samples. Furthermore, MED-Amp can be integrated with existing assays, such as mutation detection in circulating tumor cells ^30^ and next-generation sequencing of samples with limited DNA ^13^. These qualities make MED-Amp a potentially versatile tool that could be easily integrated into clinical studies.

## Materials & Methods

### Reagents and Materials

Horizon *KRAS* Gene-Specific Multiplex Reference Standard (Catalog ID HD780, Horizon Discovery), Qiagen MiniElute PCR Purification (Cat. No. 28004, Qiagen), Q5® Hot Start High-Fidelity (Catalog # M0494S, New England BioLabs® Inc.), *PfuUltra* II Fusion HS (Catalog # 600670, Agilent Technologies), Platinum™ SuperFi™ (ThermoFisher Scientific), UltraPure™ DNase/RNase-Free distilled water (Cat. No. 10977023, ThermoFisher Scientific), TaqMan® Genotyping Master Mix (Cat. No. 4371355, ThermoFisher Scientific)

### Preparation of *KRAS* genomic standards for assay benchmarking

Genomic standards were created with controlled mutant *KRAS* content by diluting Horizon *KRAS* Gene-Specific Multiplex Reference Standard with BxPC3 cell line genomic DNA. Standards were made to cover allelic frequencies ranging from 10.33% to 0.02% and total DNA content ranging from 50 ng to 5 ng. Standards were prepared in UltraPure™ DNase/RNase-Free distilled water.

### Preamplification using ddPCR

A maximum of 8 μL of DNA template was used for PCR reactions using one of three high-fidelity polymerases: Q5® Hot Start High-Fidelity, *PfuUltra* II Fusion HS, and Platinum™ SuperFi™. The RainDance Source digital PCR system was used to partition the reaction mix into approximately 5 million droplets, each five picoliters in volume (Figure 1). PCR strips containing emulsified droplets were run in a thermocycler for 9 cycles of preamplification (PCR protocols available in Supplementary Information). The droplet suspension was de-emulsified using droplet destabilizer, and residual carrier oil removed. The PCR product was processed using the Qiagen MiniElute PCR Purification kit as specified, with an additional 5 minute incubation at 35°C prior to final spin down. The elution volume for all samples was 10 μL, and samples were stored at −20°C until further use.

### ddPCR detection of preamplified template

5 μL of each sample was added to 12.5 μL of TaqMan Genotyping Master Mix, 0.9 μL of 25 μM primers, 1 μL of droplet stabilizer, 0.15 μL of 12.5 μM probes for *KRAS* wild-type/G12C/G12D/G12R/G12V (probe sequences available in Supplementary Information), and water to a final volume of 25 μL. The RainDance Source digital PCR system was used again to partition the reaction mix, and emulsified droplets were processed in a thermocycler with a 10 min annealing step at 95°C, followed by 45 cycles of 95°C for 15 s and 60°C for 1 min, concluding with a 10 min extension at 98°C. Droplets were processed using the RainDance Sense digital PCR system, and resulting populations were gated using RainDance Analyst II™ software.

### Patient Characteristics

Seventeen plasma samples were collected after Institutional Review Board approval (HUM25339) at University of Michigan and under compliance with HIPPA guidelines. Ten samples were collected from patients undergoing routine colonoscopy. Seven samples were collected from metastatic PDA patients. Eight patients were female, nine male. The average age was 54 for healthy controls and 66 for PDA patients (Table 1). Two patients had prior, or ongoing, chemotherapy and radiation treatment.

### Patient plasma collection

Patient blood samples were drawn in either Streck or EDTA tubes and were processed within 30 minutes of collection. Samples were centrifuged for 10 minutes at 820 xg at 4°C. The plasma supernatant was extracted via pipette and aliquoted in 1 mL volumes in 1.5 mL Eppendorf tubes. Plasma samples underwent a second spin at 16,000 xg for 10 minutes at 4°C. Samples were stored at −80°C until further processing. Matching buffy coat was also collected and stored at −80°C.

### Patient plasma isolation

200 μL of Proteinase K was added to all plasma samples, which were then processed using the QIAmp® Circulating Nucleic Acid kit as specified. Purified DNA samples were eluted in 150 μL Buffer AVE (RNase-free water with 0.04% sodium azide) and stored at −80°C until further use. Samples were concentrated using a standard ethanol precipitation protocol. Briefly, 0.1 vol of sodium acetate, 2.5 vol of 100% ethanol, and 0.06 μg/μL of glycogen was added to each DNA aliquot and stored overnight at −80°C. Aliquots were centrifuged at 12,000 x g at 4°C for 30 min. The supernatant was decanted and the pellet was rinsed once with ice cold 70% ethanol. The supernatant was decanted, and samples were left to air dry before resuspension in 10 μL of UltraPure™ DNAase/RNAase-free distilled water. Samples were quantified using the Qubit™ 3.0 Fluorometer High Sensitivity Kit. Final sample DNA concentrations ranged from 0.16 to 2.3 ng/μL.

### Statistical analysis

Wilcoxon rank-sum or Kruskal-Wallis tests were used to compare differences between groups. The Conover test with holm p-value correction was used for posthoc analysis. Data analysis was performed using R 3.4.4, a multi-platform open-source language and software for statistical computing ^31^.

## ASSOCIATED CONTENT

### Supporting Information

Supplementary tables & figures (pdf)

SI Table 1 *KRAS* primer and probe sequences.

SI Table 2 Sensitivity measurements for each high-fidelity polymerase tested by input *KRAS* copy number and total allelic frequency.

SI Table 3 Sensitivity measurement for plasma spike samples for each allelic frequency tested.

SI Table 4 Patient Characteristics

SI Figure 1 Representative MED-Amp results for non-template controls for each of the high-fidelity polymerases tested.

SI Figure 2 Mean cell-free DNA concentration in non-PDA controls versus metastatic PDA samples. Measured allelic frequency for controls and PDA patient samples.

SI Figure 3 Confirmation of *KRAS* mutant signal in non-PDA controls S01, and S03 via ddPCR.

## Author Contributions

EDP designed and performed experiments, analyzed data and wrote the manuscript. DBZ designed and performed experiments, analyzed data, and identified patients for study recruitment. RWC designed and performed experiments. EQ performed experiments. SLM performed experiments, consented, and collected patient samples. HC, and KS consented and collected patient samples. DS provided patient specimens.

## ACKNOWLEDGMENT

This study was financially supported by the National Institutes of Health (CA177857; DK088945; CA016672), Doris Duke Clinical Scholar Award, CPRIT Rising Stars Award (RR160022), Andrew Sabin Family Foundation, MDACC Physician Scientist Program, and the UT Stars Award. EDP and DBZ were partially supported by an NIH T32 Training Grant (T32 CA 9676-22) through the University of Michigan Cancer Biology Program.

