## Supporting Information for "Multiplex Enrichment and Detection of Rare KRAS Mutations in Liquid Biopsy Samples using Digital Droplet Pre-Amplification"

**Affiliations**
[1] Ahmed Center for Pancreatic Cancer Research, Department of Gastroenterology, Hepatology and Nutrition, University of Texas MD Anderson Cancer Center, Houston, TX, [2] Division of Hepatobiliary Surgery, Department of Surgery Comprehensive Cancer Center, University of Michigan, Ann Arbor, MI

* Corresponding author:
Erica D. Pratt
University of Minnesota Twin Cities
420 Washington Ave SE
MCB 7-194
Minnesota, MN 55455


Table of Contents

| | |
|---|---|
| Table S-1 | *KRAS* primer and probe sequences |
| Table S-2 | Sensitivity measurements for each high-fidelity polymerase tested by input *KRAS* copy number and total allelic frequency. |
| Table S-3 | Sensitivity measurement for plasma spike samples for each allelic frequency tested |
| Table S-4 | Patient characteristics |
| Figure S-1 | Representative MED-Amp results for non-template controls for each of the high-fidelity polymerases tested |
| Figure S-2 | Mean cell-free DNA concentration in non-PDA controls versus metastatic PDA samples. Measured allelic frequency for controls and PDA patient samples |
| Figure S-3 | Confirmation of *KRAS* mutant signal in non-PDA controls S01, and S03 via ddPCR |

**Table S-1.** *KRAS* primer and probe sequences

| ***KRAS* Primers** | |
|---|---|
| Forward | ATTATAAGGCCTGCTGAAAATGACT |
| Reverse | TCTGAATTAGCTGTATCGTCAAGG |

| | ***KRAS* Probes** | |
|---|---|---|
| **Target** | **Sequence** | **Reporter / Quencher** |
| WT | TTGGAGCTGGTGGCGT | VIC / MGBNFQ |
| p.G12C | TGGAGCTTGTGGCGT | FAM / MGBNFQ |
| p.G12D | TGGAGCTGATGGCGT | FAM / MGBNFQ |
| p.G12R | TGGAGCTCGTGGCGT | FAM / MGBNFQ |
| p.G12V | TGGAGCTGTTGGCGT | FAM / MGBNFQ |

MGBNFQ: minor groove binder non-fluorescent quencher

**Table S-2.** Sensitivity measurements for each high-fidelity polymerase tested by input *KRAS* copy number and total allelic frequency.

| | | Sensitivity | | |
|---|---|---|---|---|
| Allelic Frequency (%) | Total DNA Input (ng) | Q5 | PFU2 | SuperFi |
| 0.02 | 50 | 71.4 | -- | -- |
| 0.04 | 50 | 100 | -- | -- |
| 0.05 | 20 | 100 | 60 | 80 |
| 0.08 | 50 | 100 | -- | -- |
| 0.11 | 10 | 63.7 | -- | 60 |
| 0.21 | 5 | 86 | 100 | 100 |
| 0.42 | 5/10 | 93 | 100 | 100 |
| 0.49 | 20 | 100 | 100 | 100 |
| 0.83 | 5 | 91 | -- | -- |
| 0.97 | 10 | 100 | 100 | 100 |
| 4.55 | 20 | 100 | 100 | 100 |
| 8.71 | 10 | 100 | 100 | 100 |

| | Sensitivity | | |
|---|---|---|---|
| Mutant Copies | Q5 | PFU2 | SuperFi |
| 3 | 75 | 60 | 70 |
| 6 | 95 | -- | -- |
| 12 | 96 | 100 | 100 |
| 28 | 100 | 100 | 100 |
| 272 | 100 | 100 | 100 |

**Table S-3.** Sensitivity measurement for plasma spike samples for each allelic frequency tested

| Mutant Allelic Fraction | Proportion Detected | Average No. of Droplets |
|---|---|---|
| 8.71% | 100% | 560 |
| 0.97% | 66% | 39 |
| 0.49% | 100% | 84 |
| 0.21% | 100% | 206 |
| 0.05% | 33% | 24 |

**Table S-4**. Patient characteristics

| Characteristics | PDA n=7 | | Benign n=10 | |
|---|---|---|---|---|
| Age, median (range), yrs | 66 | (55-74) | 51.5 | (46-83) |
| Gender, no. (%) | | | | |
| Female | 4 | (57.1) | 7 | (70) |
| Male | 2 | (28.6) | 3 | (30) |
| Tumor Location, no. (%) | | | | |
| Head | 4 | (57.1) | -- | |
| Tail | 2 | (28.6) | -- | |
| Body | 1 | (14.3) | -- | |
| Prior Therapy, no. (%) | | | | |
| Chemotherapy | 2 | (28.6) | -- | |
| Radiation | 1 | (14.3) | -- | |
| None | 5 | (71.4) | -- | |

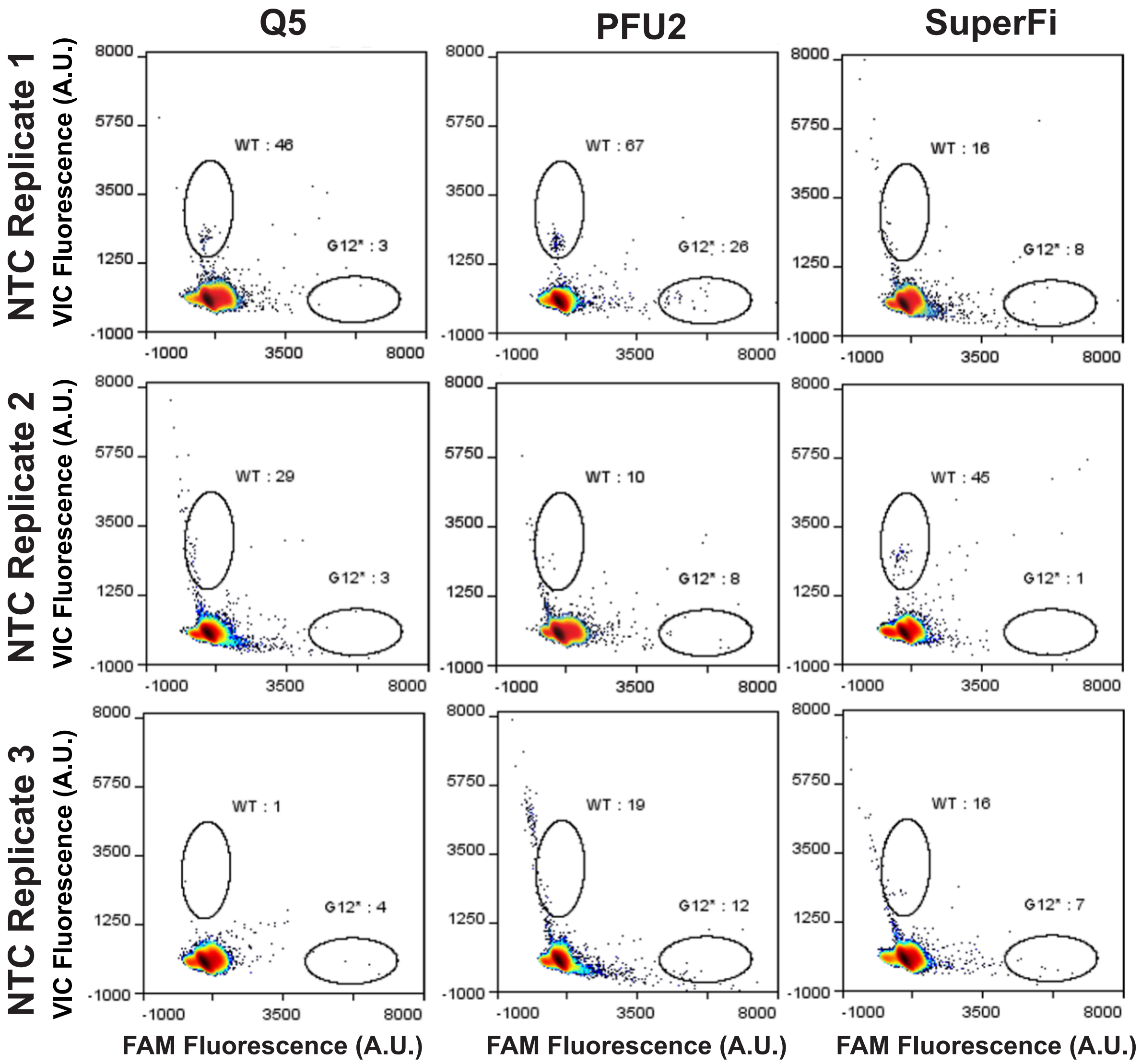


**Figure S-1.** Representative PI-AMP results for non-template controls (NTC) for each of the high-fidelity polymerases tested (Q5, PFU2, and SuperFi).

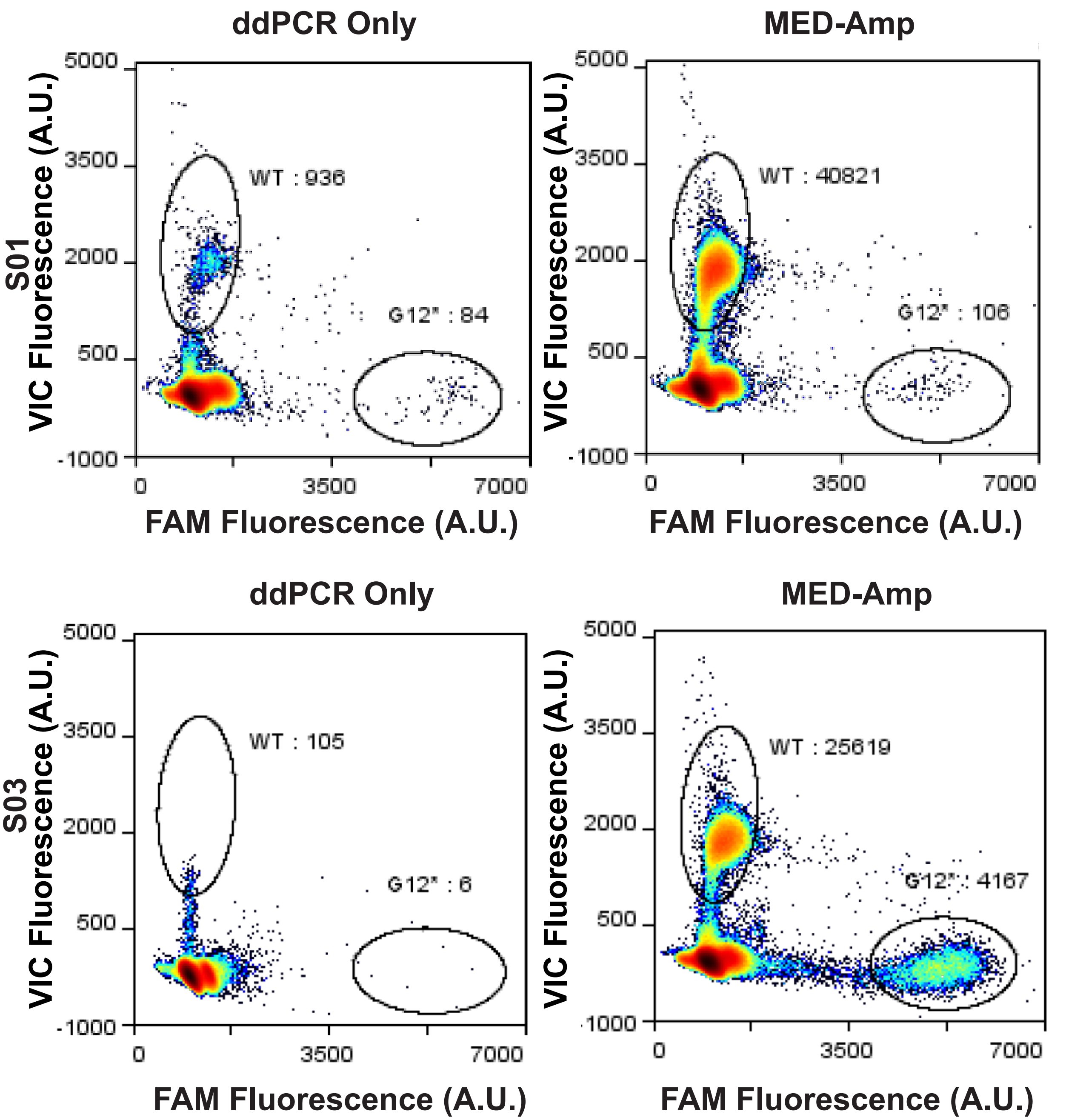


**Figure S-3.** Confirmation of *KRAS* mutant signal in non-PDA controls S01, and S03 via ddPCR. Comparison of *KRAS* signal with and without PI-AMP confirms mutant signal in the original plasma sample (>5 droplets defined as positive signal).

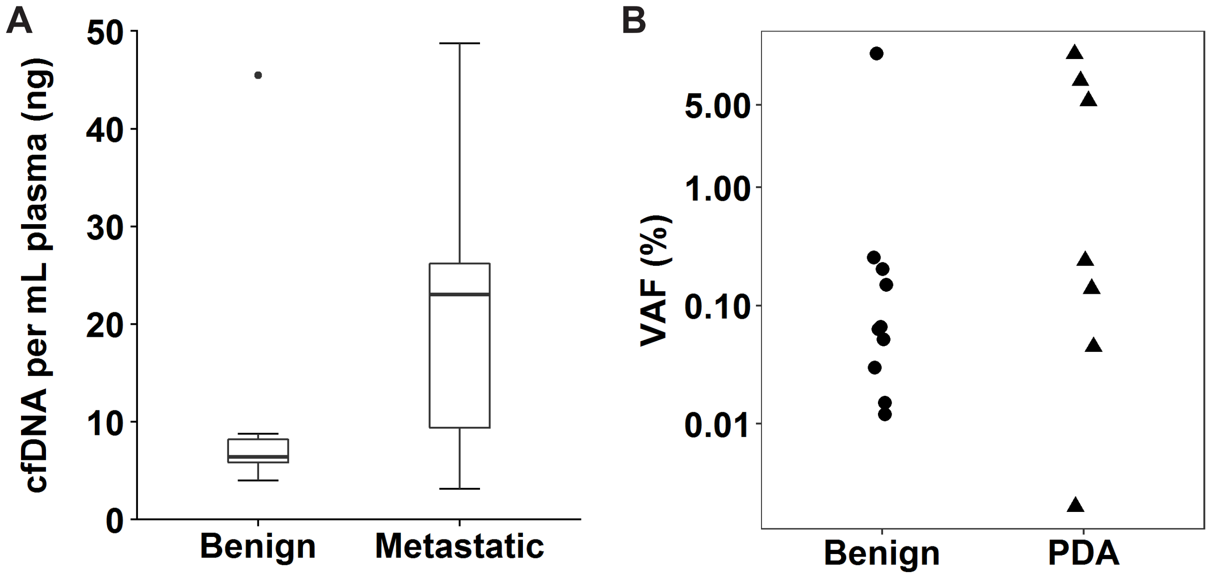


**Figure S-2.** A) Mean cell-free DNA (cfDNA) concentration in non-PDA controls versus metastatic PDA samples. B) Measured allelic frequency for controls (n=10) and PDA patient samples (n=7).
